# Connectivity and function are coupled across cognitive domains throughout the brain

**DOI:** 10.1101/2025.07.09.663763

**Authors:** Kelly J. Hiersche, Zeynep M. Saygin, David E. Osher

## Abstract

Decades of neuroimaging have revealed that the functional organization of the brain is roughly consistent across individuals and at rest it is resembles group-level task-evoked networks. A fundamental assumption in the field is that the functional specialization of a brain region arises from its connections to the rest of the brain, but limitations in the amount of data that can be feasibly collected in a single individual, leaves open the question: Is the association between task activation and connectivity consistent across the brain and many cognitive tasks? To answer this question, we fit ridge regressions models to activation maps from 33 cognitive domains (generated with NeuroQuery) using resting-state functional connectivity data from the Human Connectome Project as the predictor. We examine how well functional connectivity fits activation and find that all regions and all cognitive domains have a very robust relationship between brain activity and connectivity. The tightest relationship exists for higher-order, domain-general cognitive functions. These results support the claim that connectivity is a general organizational principle of brain function by comprehensively testing this relationship in a large sample of individuals for a broad range of cognitive domains and provide a reference for future studies engaging in individualized predictive models.

## Introduction

Decades of neuroimaging have revealed that the functional organization of the brain is roughly consistent across individuals, with e.g. ventrotemporal cortex engaged during visual perception (Grill-Spector & Weiner, 2014; Kanwisher, 2010), frontotemporal cortex engaged for linguistic processing (Fedorenko et al., 2024; Malik-Moraleda et al., 2021), and medial temporal cortex for episodic memory (Eichenbaum et al., 2012; Squire et al., 2004). Further, the functional organization of the brain at rest is also roughly consistent across individuals (Ji et al., 2019), and these resting-state networks qualitatively resemble group-level task-evoked networks (Molloy & Osher, 2023; Yeo et al., 2011). A fundamental assumption in the field is that the functional specialization of a brain region arises from its connections to the rest of the brain (Passingham et al., 2002), but is the association between task activation and connectivity consistent across the brain and many cognitive tasks?

A large body of work has been dedicated to empirically testing the relationship between connectivity and function, to predict individualized activation on a voxelwise scale. This has been done with structural connectivity in temporal cortex (Saygin et al., 2012) and across the brain (Osher et al., 2016), as well as with functional connectivity, such that an individual’s connectivity data can be used to predict their individual pattern of activation (‘connectivity fingerprint’ modeling; (Bernstein-Eliav & Tavor, 2024; Mennes et al., 2010; Molloy et al., 2024; Osher et al., 2019; Tavor et al., 2016; Tobyne et al., 2018). More recent work has demonstrated generalizability predicting individualized activation across tasks, ages, and scanner sites (Tik et al., 2023). However, despite this body of work dedicated to understanding the relationship between the idiosyncrasies of connectivity and task activation, their relationship across the broader landscape of cognition is still unclear. Does the prediction ceiling differ across different domains of processing? For example, perhaps unimodal, lower-level sensory activation is better fit by connectivity, since these regions show strong relationships between structural connectivity and function (Valk et al., 2022). Further, lower-level cognitive tasks (e.g., vision, motor) also have lower interindividual variability (Mueller et al., 2013) and develop their functionality quite early in life without substantial environmental input (Bradley & Mistretta, 1975), being instead determined by thalamic connections and under strong genetic influence (Felleman & Van Essen, 1991; Hamasaki et al., 2004; O’Leary et al., 2007). Alternatively, perhaps higher-level cognitive functions are those with the tightest link between functional activation and connectivity because they require years of co-activation and organization, leading to stronger coupling of these networks (Kelly & Castellanos, 2014; Tavoni et al., 2017). Higher-level functions also integrate information across many regions (Cohen & D’Esposito, 2016; Khodaei et al., 2023; Shine et al., 2016). Further, some cognitive functions are lateralized, showing greater activation in one hemisphere (e.g., language is lateralized to the left whereas face and social processing are lateralized to the right, (McCarthy et al., 1997; Ojemann, 1991; Rajimehr et al., 2022; Rossion et al., 2012). If repeated co-activations of regions drive tighter relationship between resting-state connectivity and functional activation, we may expect connectivity and function to be more tightly linked in the dominant hemisphere for lateralized skills but show comparable associations across hemispheres for functions that recruit bilaterally.

The goal of this study is study is to provide a bird’s eye view of the relationship between functional connectivity and functional activation across the entire brain for a broad sample of cognitive processing. To do this, we leveraged a large, publicly available database (Human Connectome Project, HCP, (Van Essen et al., 2013)) and NeuroQuery (Dockès et al., 2020), an online metanalytic tool that predicts brain maps (activation maps) based on semantic similarity among search queries and terms reliably associated with brain regions. We implemented ridge regression modeling to examine the association between functional connectivity and task-activation to a broad range of cognitive domains across the entire brain. We find that all regions and all cognitive domains have a very robust relationship between brain activity and connectivity, with the tightest relationship for higher-order, domain-general cognitive functions. These results support the claim that connectivity is a general organizational principle of brain function by comprehensively testing this relationship in a large sample of individuals for a broad range of cognitive domains and provide a reference for future studies engaging in individualized predictive models.

## Results

Functional activation was successfully modeled by resting state connectivity across all domains and all regions (see two example fitted vs expected activation maps in **Figure 1A**, see **Figure S1** for all models fits), and each model’s fit was higher than all random permutations. Model fits across all 2706 models ranged from 0.35 to 0.9997, with a strong left skew, such that 1292 models had a fit above 0.9 whereas only 26 models had a fit under 0.5, and the median fit across all models was 0.89. When examining median model fit across all regions within a task, we see that model fit remained high for all domains. The median fit (followed by random permutation 99^th^ percentile fit) was 0.88 (0.46) for the sensory domain, 0.90 (0.46) for somatosensory, 0.88 (0.46) for language, 0.88 (0.46) for social cognition, 0.89 (0.53) for decision making, 0.91 (0.56) for memory, and 0.91 (0.51) for executive functioning (see **Figure 1B** for fit by task). Model fit was also high when examining median model fit for all tasks within a lobe: 0.91 (0.77) for occipital, 0.85 (0.14) for temporal, 0.88 (0.31) for parietal, 0.90 for frontal (0.35), 0.95 (0.30) for cingulate, and 0.93 (0.19) for subcortical (see **Figure 2**).

**Figure 1:**
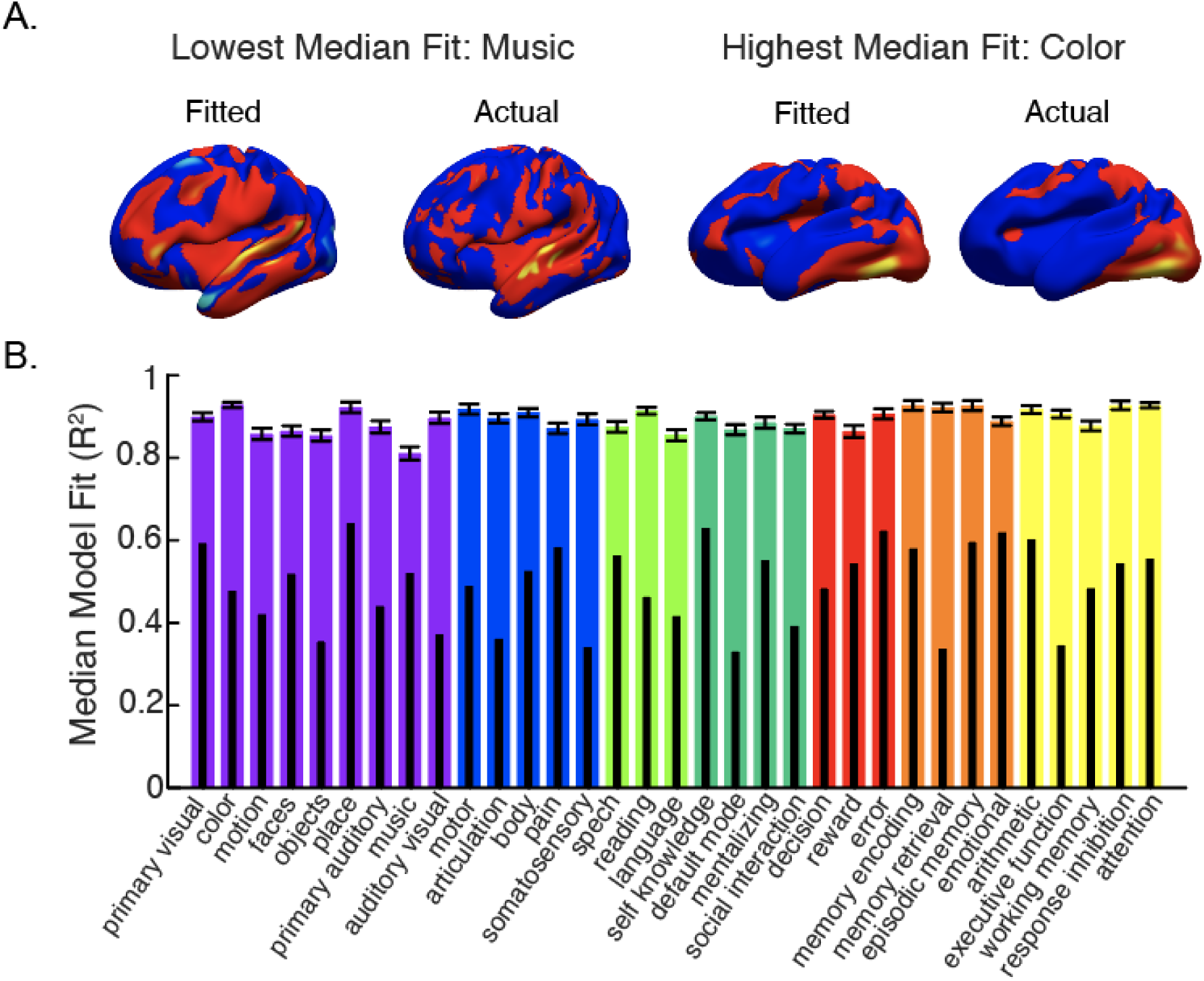
A. Expected activation maps (from NeuroQuery, left) and connectivity-based fitted activation (right) for two sample cognitive domains. B. Large colored bars show median fit (with standard error bars) for each domain across regions. Subset black bars show the 99^th^ percentile performance of the permuted models for each domain. Sensory is in purple, somatosensory is in blue, language is in green, social is in blue-green, decision making is in red, memory is in orange, and executive functioning is in yellow.

**Figure 2.**
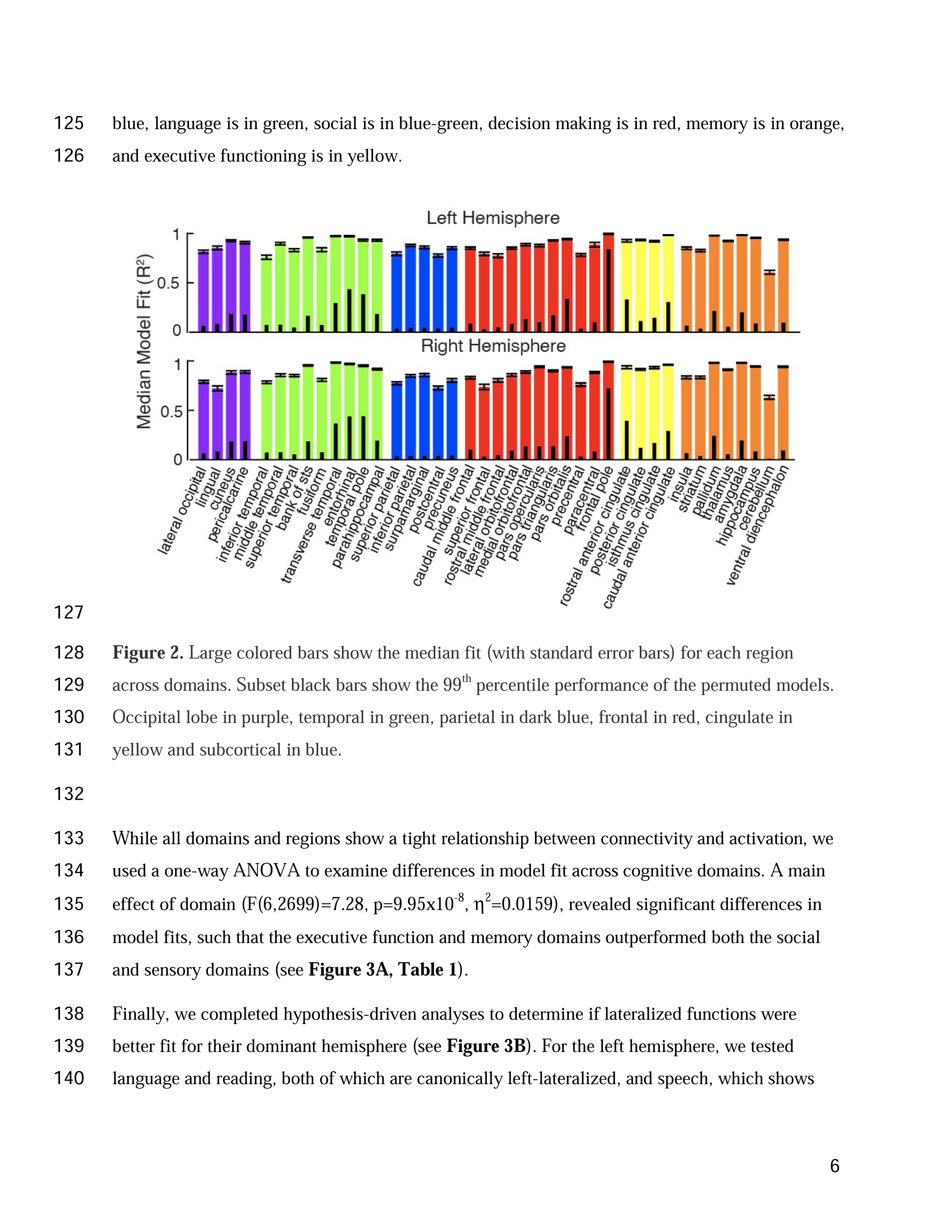
Large colored bars show the median fit (with standard error bars) for each region across domains. Subset black bars show the 99^th^ percentile performance of the permuted models. Occipital lobe in purple, temporal in green, parietal in dark blue, frontal in red, cingulate in yellow and subcortical in blue.

While all domains and regions show a tight relationship between connectivity and activation, we used a one-way ANOVA to examine differences in model fit across cognitive domains. A main effect of domain (F(6,2699)=7.28, p=9.95×10^-8^, η^2^=0.0159), revealed significant differences in model fits, such that the executive function and memory domains outperformed both the social and sensory domains (see **Figure 3A**, **Table 1**).

**Figure 3:**
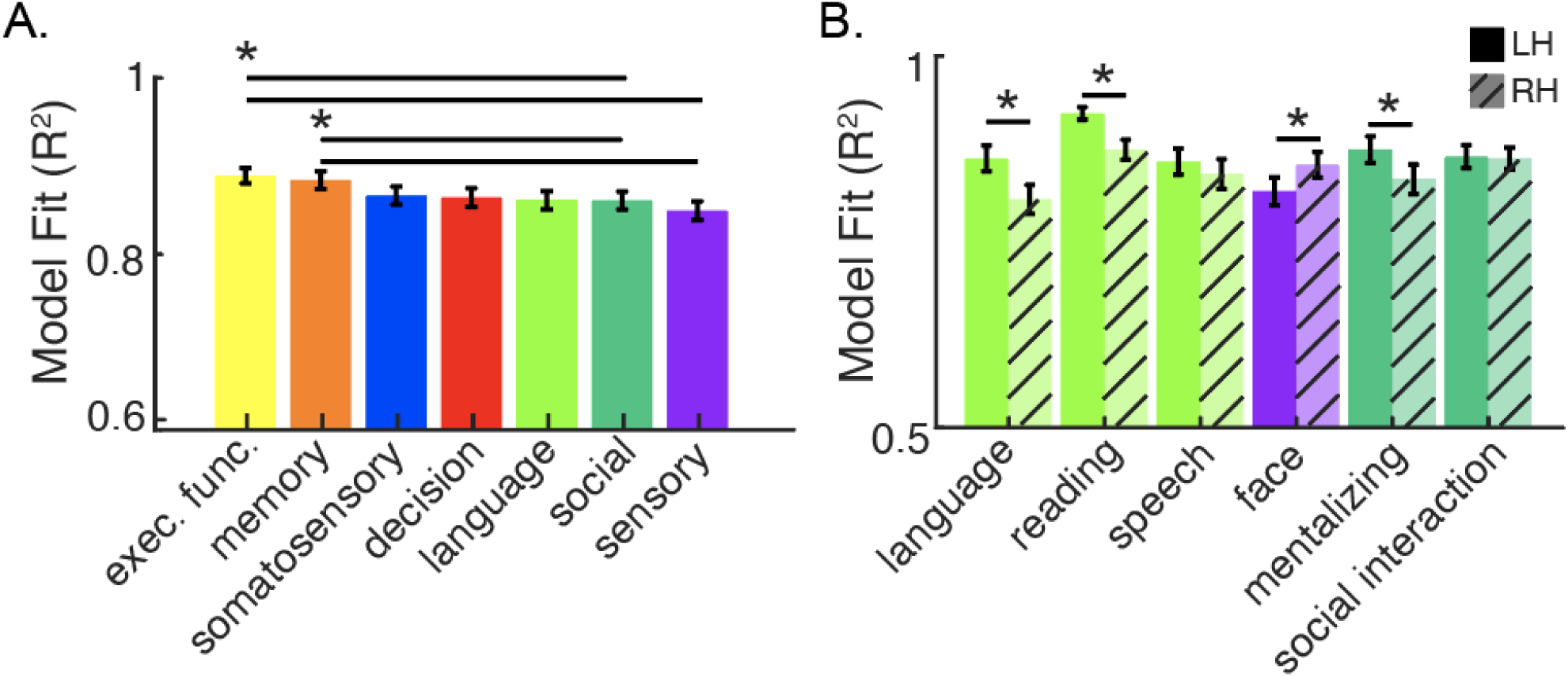
Results of one-way ANOVA and test of lateralized functions. A. Main effect of category with mean model fit with standard error bars. *p<0.05, surviving Bonferroni-Holm multiple comparison corrections for 21 comparisons. B. Hemispheric differences for lateralized functions. Mean model fit with standard error bars. *p<0.05, Bonferroni-Holm corrected for 6 comparisons.

**Table 1:** Results of post-hoc two-tailed, independent samples t-tests to compare model fit across all 7 cognitive domains. Uncorrected p-values listed and Cohen’s d effect sizes. **p<0.05 after Bonferroni-Holm multiple comparisons correction for 21 tests.

| Category | Category | *t* (df) | *p* | *d* |
| --- | --- | --- | --- | --- |
| decision making | executive function | -2.62 (453) | 0.0090 | -0.215 |
| decision making | language | 0.43 (490) | 0.66 | 0.039 |
| decision making | memory | -2.32 (520) | 0.02 | -0.197 |
| decision making | sensory | 1.73 (438) | 0.08 | 0.126 |
| decision making | social | 0.58 (515) | 0.56 | 0.049 |
| decision making | somatosensory | -0.18 (507) | 0.86 | -0.015 |
| executive function | language | 3.17 (460) | 0.0020** | 0.260 |
| executive function | memory | 0.079 (653) | 0.94 | 0.006 |
| executive function | sensory | 5.76 (997) | 1.10x10<sup>-8**</sup> | 0.344 |
| executive function | social | 3.67 (660) | 2.64x10<sup>-4**</sup> | 0.274 |
| executive function | somatosensory | 2.89 (802) | 0.0040 | 0.202 |
| language | memory | -2.83 (527) | 0.0050 | -0.239 |
| language | sensory | 1.23 (447) | 0.22 | 0.089 |
| language | social | 0.11 (521) | 0.91 | 0.009 |
| language | somatosensory | -0.67 (516) | 0.50 | -0.054 |
| memory | sensory | 4.88 (671) | 1.31x10<sup>-6**</sup> | 0.319 |
| memory | social | 3.21 (654) | 0.0010** | 0.251 |
| memory | somatosensory | 2.49 (702) | 0.010 | 0.184 |
| sensory | social | -1.24 (683) | 0.22 | -0.081 |
| sensory | somatosensory | -2.317 (895) | 0.021 | -0.141 |
| social | somatosensory | -0.87(707) | 0.39 | -0.064 |

Finally, we completed hypothesis-driven analyses to determine if lateralized functions were better fit for their dominant hemisphere (see **Figure 3B**). For the left hemisphere, we tested language and reading, both of which are canonically left-lateralized, and speech, which shows bilateral activation, and found that connectivity significantly better fit functional activation in the LH for language (t(40)=3.32, p=0.0019, d=0.45) and reading (t(40)=6.08, p=3.64×10^-7,^ d=0.63), whereas fit did not differ across the hemispheres for speech (t(40)=1.28, p=0.21, d=0.12). We also tested mentalizing and face perception, which show right lateralized activation, and social interaction, which elicits bilateral activation. As expected, face perception was better fit in the RH (t(40)=-2.83, p=0.0072, d=-0.31) and social interaction was equally well fit across the hemispheres (t(40)=0.30, p=0.76, d=0.025). Surprisingly, we found that mentalizing was better fit in the LH (t(40)=3.01, p=0.0.0045, d=0.28), in contradiction with our hypothesis that functions would be better fit for their dominant hemispheres (mentalizing, similar to other social cognitive skills, is lateralized to the RH (Gupta et al., 2025; Rajimehr et al., 2022, 2022)). All comparisons survived correction for 6 multiple comparisons.

## Discussion

Here we provide substantial evidence that connectivity is a general organizational principle of brain function across the entire brain. Voxelwise functional activation for each region of the brain is well explained by its connectivity to other regions, and this is true across a broad array of cognitive domains and across the entire brain. Our analyses also reveal differences in model fits across cognitive domains and hemispheres, which we discuss below.

This study provides a bird’s eye view of the functional connectivity and functional activation relationships across the entire brain for a broad sample of cognitive processing. Even with access to large-scale databases with many tasks and subjects, connectivity-function relationships have only ever been tested in as many as 7 domains at a time (Tavor et al., 2016). This work demonstrates that connectivity-activation relationships are robust enough to emerge even when the data are representative of general brain activation and not explicitly measured in the same individuals. This provides quantitative evidence for a relationship that has been largely assumed in the past and tested in only a small sample of potential cognitive domains. Further, we show that connectivity can explain expected activation in regions that are important for a task (e.g., language activation in temporal regions), but also regions that are not involved in a task (occipital activation in speech). Given that the nuances of activation across all regions can modeled, regardless of a region’s activation in a task, this paper provides a noise ceiling for the expected fit of connectivity and function for researchers involved in individualized predictive approaches. For example, our work suggests that the cerebellum is not well modeled when treated as a single anatomical region. Therefore, parcellations of the cerebellum, allowing for different models for different subregions, may be important for individuals attempting to predict cerebellar function based on its connectivity. Further, given that this model allows us to examine connectivity-function relationship across the whole brain, this model could be altered by ‘lesioning’ particular regions (either typically involved or not in a task), to make predictions about expected changes in functional activation. Additionally, these results provide a baseline for what might be considered the norm relationship between connectivity and function, allowing future work on individual differences to compare how an individual may vary from this average expected relationship. Finally, our study provides a reference for connectivity-function relationships in a young, healthy sample, and a framework for assess these relationships in other populations. Given evidence suggests that connectivity patterns may be altered in aging and disease (Farràs-Permanyer et al., 2019; Guimarães et al., 2017; Sala-Llonch et al., 2015; Zhang et al., 2010), future work can apply this approach to characterize alterations in the connectivity-function relationship in aging populations, developmental samples, and disease.

Our planned analyses showed that lateralized functional activation (e.g. for language and faces) was better explained by connectivity in the dominant hemisphere, whereas bilateral functions within similar domains were equally well fit across the hemispheres. Lateralization of function is an interesting nuance of human cognition, and it indeed makes sense that connectivity patterns mimic this laterality. Certain functional lateralities develop quite early in development and may be driven by lateralized connectivity patterns that are genetically determined. For example, the language network tends to be left-lateralized by age 3, although this specialization and related lateralization continues to develop through early childhood (Hiersche et al., 2024; Olulade et al., 2020). Perhaps this early-developing functional laterality is biased or driven by connectivity (e.g. (Li et al., 2020; Saygin et al., 2016)), suggesting that the tight link between connectivity and function that we observe here with adults may reflect early developmental biases. However, we also observed stronger fits with connectivity for mentalizing on the LH, despite greater RH activation. This result suggests that there may also be other factors that drive certain connectivity-function relationships, such as ongoing maturation and continued synchronization and strengthening of these networks, resulting in greater engagement of the LH in aspects of mentalizing that may require more effort and experience (Doricchi et al., 2022; Sun et al., 2023).

In line with this idea, executive function and memory domains which included arithmetic, attention, working memory, response inhibition, attention, episodic memory, and memory encoding and retrieval were the best fit by connectivity and significantly outperformed social and sensory domains. Executive function skills (and the networks that support them) show prolonged maturation requiring many years to develop (Best & Miller, 2010; Gao et al., 2015; Sherman et al., 2014). This contrasts with cognitive skills and networks that are in place very early in development. For example, sensory networks are in place in utero and mature rapidly (Eyre et al., 2021), whereas higher-order association networks are not always detectable or are weaker (Thomason et al., 2013). Thus, the stronger fit of connectivity and functional activation in these higher-order domains may reflect the ongoing strengthening of disparate nodes of multimodal cortex and their connections through repeated co-activations (e.g. through interactive specialization, (Johnson, 2011)).

By utilizing the large-scale data available through HCP and the meta-analysis tool NeuroQuery, this study examined the relationship between connectivity and function at a greater scale than any prior work. While aligning with prior showing resting-state functional connectivity is strongly related to task activations (Cole et al., 2016), this method allowed us to examine a wider range of cognitive domains, testing the coupling of connectivity and function from low-level sensory activation to complex cognitive tasks, which is not feasible in any single data set collected. However, this approach also has its limitations. The ideal case would be to have functional connectivity and task activation data in the same set of individuals, across many possible cognitive domains. Further, while the cognitive domains selected do cover a breadth of terms studied by cognitive neuroscientists, we have not covered all possible terms; although given the consistency across all tested categories, it seems likely that the association between connectivity and function is strong for other domains as well. Further, we provide the code for these analyses (see Data & Code Availability section), allowing other researchers to expand upon the chosen domains for their own work, when applicable. Also, there is the possibility that NeuroQuery utilized studies implementing connectivity-based analyses when generating its predictive maps; therefore, there is potential that the relationship between connectivity and function could be mildly inflated, though given the number of papers that contributed to NeuroQuery (over 13,000) and the much greater prevalence of task-based fMRI research, likely many studies did not use this method. Further, while we build upon the body work examining the connectivity-function relationship (for review see: (Bernstein-Eliav & Tavor, 2024)) we were unable to examine individual variability of this connectivity-function relationship across domains with this meta-analytic approach. Despite this limitation, we hope this study can be a helpful benchmark for estimating the potential of individualized predictive models across numerous domains and regions. Finally, this work does not consider or test the role of multivariate representations, which could reveal different aspects of information processing or more nuanced representations (e.g., differentiating exemplars of a particular category or domain of knowledge like cats vs. dogs, or dynamic facial expressions) with differing relationships to connectivity.

Numerous questions remain. Future work should explore the contributions of structural connectivity, which has also been used to predict individualized activation (Osher et al., 2016; Saygin et al., 2012), as well as cortical folding patterns which create subject-specific anatomical fingerprints (Duan et al., 2020). These anatomical measures will offer other potential mechanisms for complex mental function and likely interact with co-activations and functional connectivity in interesting ways (e.g (Bullmore & Sporns, 2009; Sporns, 2013)). Further, given that an individual’s functional connectivity fingerprint may be genetically driven (Hu et al., 2022), genetic mechanisms of connectivity fingerprints should be explored to understand the influence of genetic vs. experiential contributions to the link between connectivity and function. Finally given the changes to functional connectivity across the lifespan (Betzel et al., 2014), future work should examine potential shifts in connectivity-function coupling across the lifespan. Longitudinal, developmental work can be used to further test whether connectivity-function coupling is strongest for tasks requiring greater environmental experience and determine the trajectory of the relationship between function and connectivity across the brain for different cognitive domains. This study provides the foundation for these next steps, allowing future work to further characterize the mechanisms supporting brain function.

## Methods

### rs-fMRI Data

Each participant who completed all 4 runs of resting state fMRI from the Human Connectome Project (HCP; (Van Essen et al., 2013)) was included for connectivity analyses (N=1018; 546 females, 472 males). Four 15-min runs of resting state data per individual were preprocessed following the HCP minimal preprocessing pipeline (Glasser et al., 2013). Additional preprocessing of the resting state data included denoising with aCompCor (Behzadi et al., 2007), nuisance regression using the top 5 temporal principal components in white matter and CSF separately, and surface smoothed using a gaussian kernel with σ=3mm. Functional connectivity was computed for each subject, per run, as the Pearson correlation coefficient between each vertex and the mean timecourse of each region in the Desikan-Killiany atlas (Desikan et al., 2006). Data were Fisher-transformed and averaged across runs, and a final group average connectivity matrix was computed as the mean across all subjects. This matrix contained the average functional connectivity of each vertex to each cortical and subcortical region, and was thus of size 91282 vertices by 89 regions. The resulting matrix supplied the covariates of each multivariate model described below.

### NeuroQuery Activation Maps

We selected 33 cognitive domains of broad interest to cognitive neuroscientists, spanning seven categories (sensory, somatosensory, language, decision making, social, memory, executive function; **see Figure 1**). Each term was entered into NeuroQuery (a tool that uses search queries to generate maps of fMRI responses using information from 7547 neuroimaging publications (Dockès et al., 2020) on 02/09/2025 or on 12/19/2024 and the resulting brain map was downloaded (these maps are available in volume space here, see data code & availability; and **Figure S2** shows activation on lateral surface for all categories). This map was examined by two experts in the field (authors Z.M.S. and D.E.O.), to confirm that expected brain regions were active for each term, to determine its category, (e.g., to determine that ‘body’ activated motor regions of the brain, placing it in the somatosensory category, and was not activating just high-level visual body regions in ventral temporal cortex) and that expected activation maps across domains were not highly overlapping (e.g., social interaction and mentalizing activate distinct regions of social processing; including faces, but not also facial expressions, as the maps were highly overlapping). Further, each map was visually compared to multiple similar search terms in Neurosynth (Yarkoni et al., 2011), a commonly used meta-analytic tool for examining activation to cognitive domains across a wide range of peer reviewed studies.

Results from NeuroQuery provide an expected activation (z statistic) for each 4 mm isotropic voxel in MNI152 space. Each map was upsampled to 2mm and then registered to the fsaverage_LR32k template surface using Connectome Workbench ((Marcus et al., 2011) https://github.com/Washington-University/workbench). These vertex-wise activation maps were the response variables of the multivariate models described below.

### Modeling Procedure and Analysis

In this study, we are assessing the goodness of fit of regression models that integrate resting-state connectivity with function (see **Figure 4** for modeling procedure). Given that NeuroQuery only provides a single meta-analytic brain map, there is no out-of-sample data to predict. Therefore, we fit the average connectivity matrix across many subjects to the NeuroQuery expected activation in each region, using a separate model for each brain region and domain (33 domains x 82 regions). For each cognitive domain, connectivity and activation maps were extracted for each region separately, such that each vertex of a region is described by its degree of activation (scalar) and its connectivity pattern to each region of the brain (vector). To fit connectivity data to activation maps, we utilized ridge regression models (Hoerl & Kennard, 1970), similar to how we have previously performed connectivity fingerprinting analyses (Molloy et al. 2024; Osher et al. 2019). First, the hyperparameter (λ) was optimized for each model, using a 5-fold cross-validation with 100 maximum number of objective function evaluations. Then that optimal lambda was used to fit a ridge regression model (Matlab R2020a, fitrlinear) to the HCP connectivity data and NeuroQuery meta-analytic map. We computed the coefficient of determination (R^2^), a commonly used within-sample measure for goodness-of-fit, which measures the percent variance in functional responses that is accounted for by each connectivity model.

**Figure 4:**
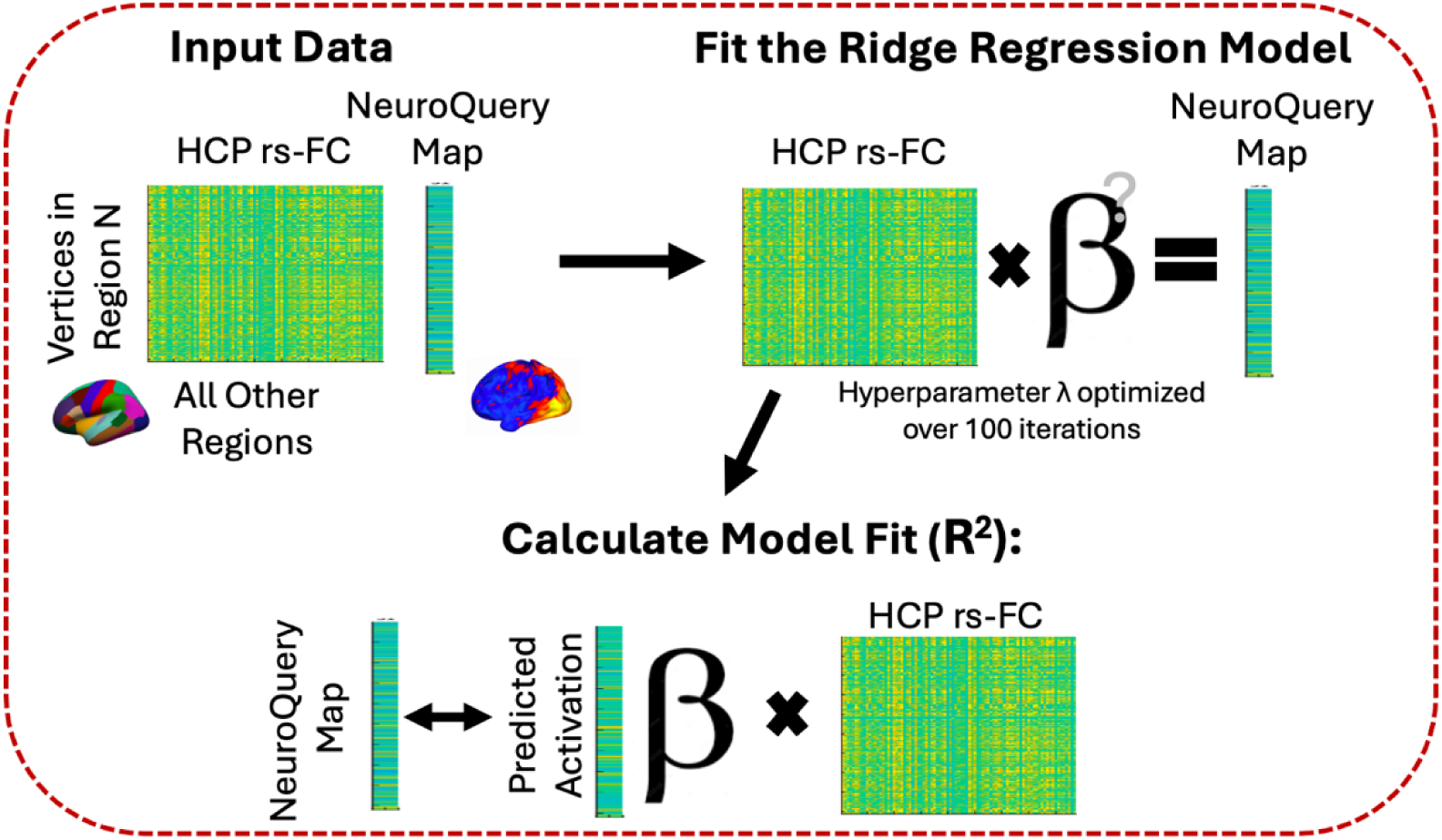
Diagram outlining the model procedures for one domain and one region. This process was repeated for all 33 domains and 82 regions.

Permutation testing determined if the relationship between functional connectivity and fitted activation for a particular region was significant. Each voxel was randomly assigned (within a region, without replacement) the functional activation of the same cognitive domain of another voxel while keeping its functional connectivity profile intact. With the permuted activation data (the dependent variable) but intact functional connectivity data (independent variables), we created 1000 new regularized ridge regression models for each region and each domain to determine the relationship between “true” connectivity and “chance” expected activation. These permutation models utilized the same hyperparameters from the “true” model. For each permutation model, we calculated R^2^ to create a distribution of model fits to test whether our “true” model fits were likely the outcome of chance or represented a real relationship between functional connectivity and activation.

### Statistical Analyses

To examine the significance of each model fit, we compared the “true” model fit (R^2^) to the distribution of ‘random’ fits calculated during permutation testing by determining the percentage of permutation models that outperformed our true model (sum permutation model R^2^ > “true” model fit divided by 1000) and calculating the 99% confidence level of the permuted models (analyzed in Matlab R2020a). Distribution of model fit across domains and regions indicated primarily skewed distributions; therefore, median model fit is reported when collapsing across domains or regions.

To determine whether all cognitive domains were equally well fit by connectivity, we used a one-way ANOVA to examine differences in model fit (R^2^) across domains. The dependent variable (R^2^) was initially negatively skewed and was transformed (arcsine of square root) to represent a more normal distribution (examined with QQ-plot), and a plot of fitted values verses residuals of the transformed data model showed a relative lack of relationship between fitted values and residuals, indicating homogeneity of variance. Two-tailed, independent sample t-tests were used for post-hoc comparisons following a significant main effect, and all reported statistics survive multiple comparison corrections for 21 comparisons. Cohen’s d effect sizes are included for all t-test comparisons.

Finally, we conducted hypothesis driven analysis (these analyses were not preregistered), based on expected laterality differences in fit for a few select cognitive domains. Paired samples, two-tailed, t-tests were used to compare fit across the hemispheres for reading, language, and speech, with the expectation that left-lateralized functions (language and reading), should show a tighter association between connectivity and function (and therefore higher model fit) in the LH compared to RH, whereas speech, which is more bilateral, should not show these differences. We also tested faces and mentalizing, which show right lateralized activation maps, and social interaction, which shows bilateral activation. All reported significant effects survive multiple comparison for the 6 tests performed.

## Data and Code Availability

Brain maps, downloaded from NeuroQuery, along with all code used for modeling and analysis are available here: https://github.com/CognitionBrainCircuitryLab/Connectivity-and-function-across-domains.

## Acknowledgements

“Data were provided [in part] by the Human Connectome Project, WU-Minn Consortium (Principal Investigators: David Van Essen and Kamil Ugurbil; 1U54MH091657) funded by the 16 NIH Institutes and Centers that support the NIH Blueprint for Neuroscience Research; and by the McDonnell Center for Systems Neuroscience at Washington University.“

---

**Figure S1:**
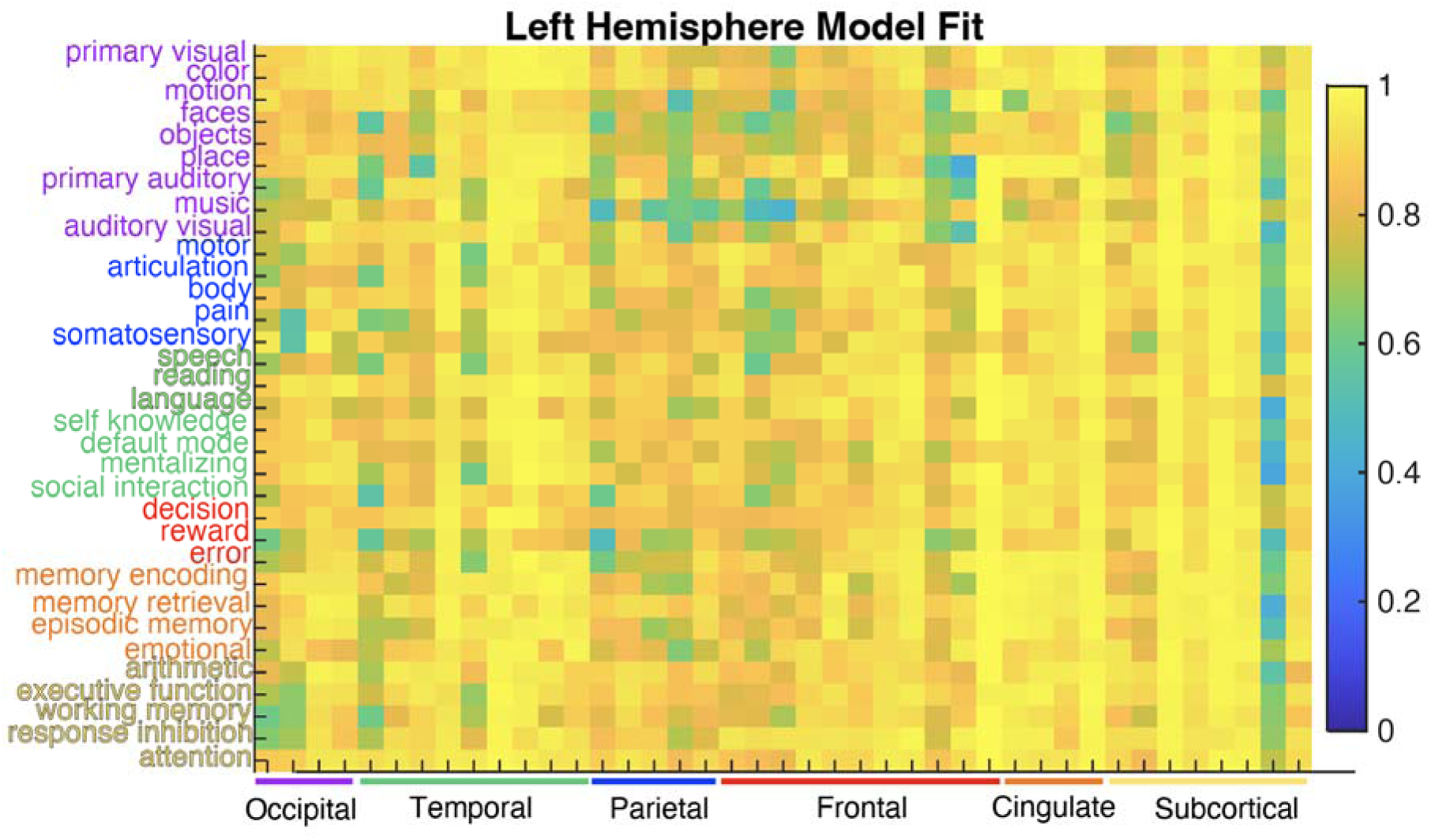
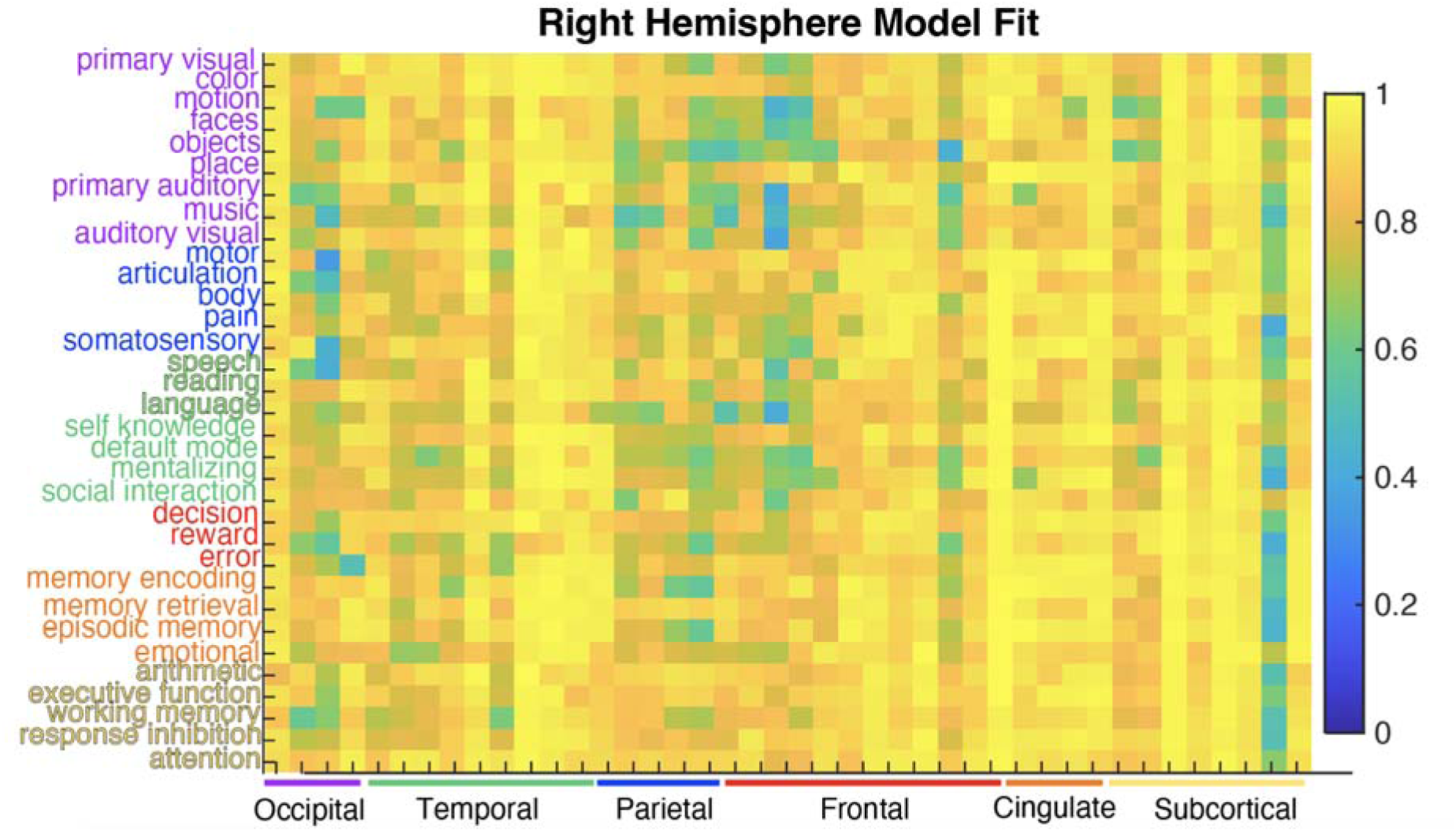
Heatmap of model fit for all domains across lobes. Regions are in the same order as presented in Figure 2.

**Figure S2:**
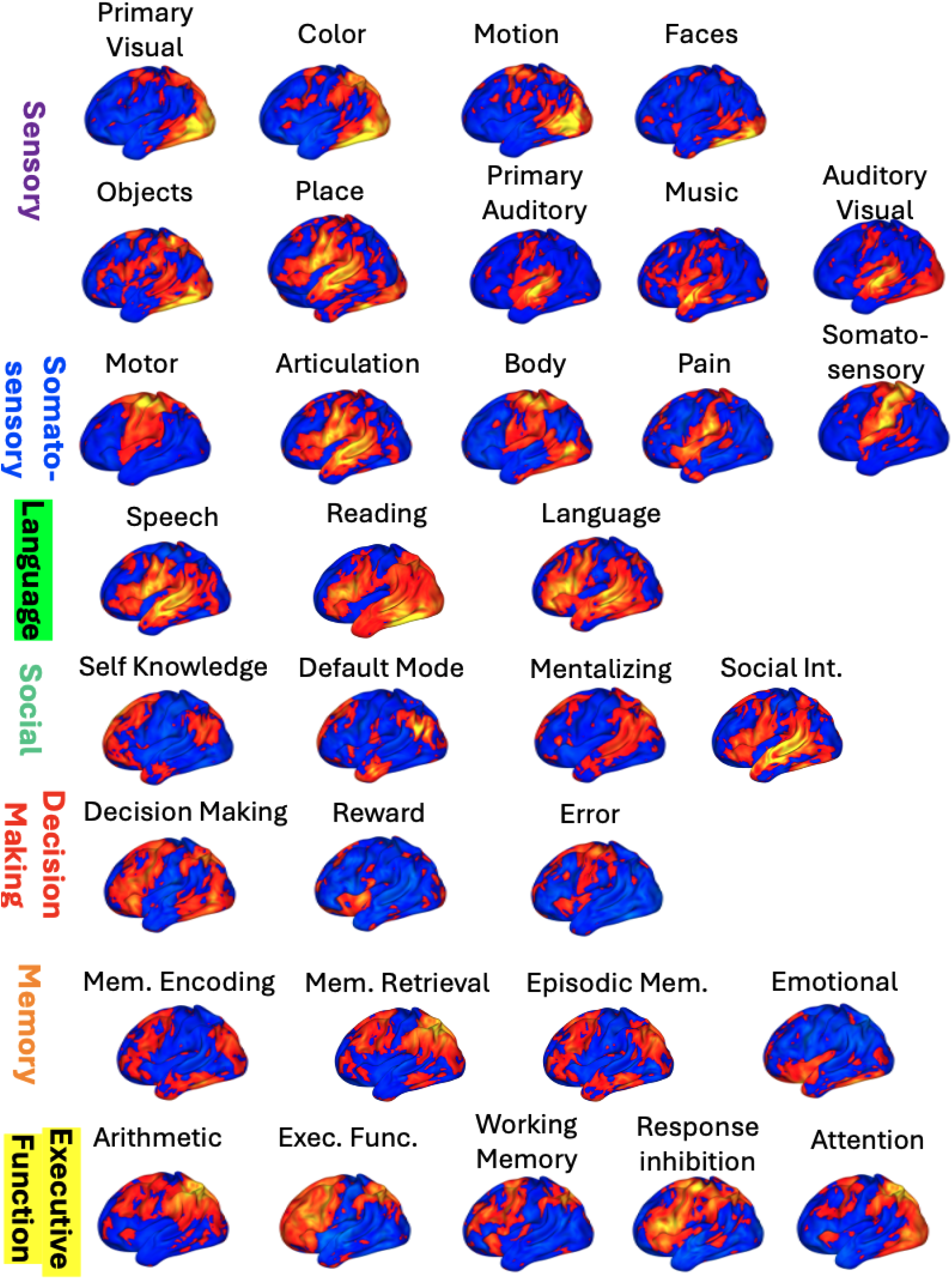
NeuroQuery expected activation maps (used as outcome variable in models) for all 33 cognitive domains on left lateral surface of FsAverage.

## References

Behzadi, Y., Restom, K., Liau, J., & Liu, T. T. (2007). A component based noise correction method (CompCor) for BOLD and perfusion based fMRI. NeuroImage, 37(1), 90–101. 10.1016/j.neuroimage.2007.04.042

Bernstein-Eliav, M., & Tavor, I. (2024). The Prediction of Brain Activity from Connectivity: Advances and Applications. The Neuroscientist, 30(3), 367–377. 10.1177/10738584221130974

Best, J. R., & Miller, P. H. (2010). A Developmental Perspective on Executive Function. Child Development, 81(6), 1641–1660. 10.1111/j.1467-8624.2010.01499.x

Betzel, R. F., Byrge, L., He, Y., Goñi, J., Zuo, X.-N., & Sporns, O. (2014). Changes in structural and functional connectivity among resting-state networks across the human lifespan. NeuroImage, 102, 345–357. 10.1016/j.neuroimage.2014.07.067

Bradley, R. M., & Mistretta, C. M. (1975). Fetal sensory receptors. Physiological Reviews, 55(3), 352–382. 10.1152/physrev.1975.55.3.352

Bullmore, E., & Sporns, O. (2009). Complex brain networks: Graph theoretical analysis of structural and functional systems. Nature Reviews Neuroscience, 10(3), 186–198. 10.1038/nrn2575

Cohen, J. R., & D’Esposito, M. (2016). The Segregation and Integration of Distinct Brain Networks and Their Relationship to Cognition. Journal of Neuroscience, 36(48), 12083–12094. 10.1523/JNEUROSCI.2965-15.2016

Cole, M. W., Ito, T., Bassett, D. S., & Schultz, D. H. (2016). Activity flow over resting-state networks shapes cognitive task activations. Nature Neuroscience, 19(12), 1718–1726. 10.1038/nn.4406

Desikan, R. S., Ségonne, F., Fischl, B., Quinn, B. T., Dickerson, B. C., Blacker, D., Buckner, R. L., Dale, A. M., Maguire, R. P., Hyman, B. T., Albert, M. S., & Killiany, R. J. (2006). An automated labeling system for subdividing the human cerebral cortex on MRI scans into gyral based regions of interest. NeuroImage, 31(3), 968–980. 10.1016/j.neuroimage.2006.01.021

Dockès, J., Poldrack, R. A., Primet, R., Gözükan, H., Yarkoni, T., Suchanek, F., Thirion, B., & Varoquaux, G. (2020). NeuroQuery, comprehensive meta-analysis of human brain mapping. eLife, 9, e53385. 10.7554/eLife.53385

Doricchi, F., Lasaponara, S., Pazzaglia, M., & Silvetti, M. (2022). Left and right temporal-parietal junctions (TPJs) as “match/mismatch” hedonic machines: A unifying account of TPJ function. Physics of Life Reviews, 42, 56–92. 10.1016/j.plrev.2022.07.001

Duan, D., Xia, S., Rekik, I., Wu, Z., Wang, L., Lin, W., Gilmore, J. H., Shen, D., & Li, G. (2020). Individual identification and individual variability analysis based on cortical folding features in developing infant singletons and twins. Human Brain Mapping, 41(8), 1985–2003. 10.1002/hbm.24924

Eichenbaum, H., Sauvage, M., Fortin, N., Komorowski, R., & Lipton, P. (2012). Towards a functional organization of episodic memory in the medial temporal lobe. Neuroscience & Biobehavioral Reviews, 36(7), 1597–1608. 10.1016/j.neubiorev.2011.07.006

Eyre, M., Fitzgibbon, S. P., Ciarrusta, J., Cordero-Grande, L., Price, A. N., Poppe, T., Schuh, A., Hughes, E., O’Keeffe, C., Brandon, J., Cromb, D., Vecchiato, K., Andersson, J., Duff, E. P., Counsell, S. J., Smith, S. M., Rueckert, D., Hajnal, J. V., Arichi, T., … Edwards, A. D. (2021). The Developing Human Connectome Project: Typical and disrupted perinatal functional connectivity. Brain, 144(7), 2199–2213. 10.1093/brain/awab118

Farràs-Permanyer, L., Mancho-Fora, N., Montalà-Flaquer, M., Bartrés-Faz, D., Vaqué-Alcázar, L., Peró-Cebollero, M., & Guàrdia-Olmos, J. (2019). Age-related changes in resting-state functional connectivity in older adults. Neural Regeneration Research, 14(9), 1544. 10.4103/1673-5374.255976

Fedorenko, E., Ivanova, A. A., & Regev, T. I. (2024). The language network as a natural kind within the broader landscape of the human brain. Nature Reviews Neuroscience, 25(5), 289–312. 10.1038/s41583-024-00802-4

Felleman, D. J., & Van Essen, D. C. (1991). Distributed hierarchical processing in the primate cerebral cortex. *Cerebral Cortex (New York*, N.Y., 1(1), 1–47. 10.1093/cercor/1.1.1-a

Gao, W., Alcauter, S., Elton, A., Hernandez-Castillo, C. R., Smith, J. K., Ramirez, J., & Lin, W. (2015). Functional Network Development During the First Year: Relative Sequence and Socioeconomic Correlations. Cerebral Cortex, 25(9), 2919–2928. 10.1093/cercor/bhu088

Glasser, M. F., Sotiropoulos, S. N., Wilson, J. A., Coalson, T. S., Fischl, B., Andersson, J. L., Xu, J., Jbabdi, S., Webster, M., Polimeni, J. R., Van Essen, D. C., & Jenkinson, M. (2013). The minimal preprocessing pipelines for the Human Connectome Project. NeuroImage, 80, 105–124. 10.1016/j.neuroimage.2013.04.127

Grill-Spector, K., & Weiner, K. S. (2014). The functional architecture of the ventral temporal cortex and its role in categorization. Nature Reviews Neuroscience, 15(8), 536–548. 10.1038/nrn3747

Guimarães, R. P., Arci Santos, M. C., Dagher, A., Campos, L. S., Azevedo, P., Piovesana, L. G., De Campos, B. M., Larcher, K., Zeighami, Y., Scarparo Amato-Filho, A. C., Cendes, F., & D’Abreu, A. C. F. (2017). Pattern of Reduced Functional Connectivity and Structural Abnormalities in Parkinson’s Disease: An Exploratory Study. Frontiers in Neurology, 7. 10.3389/fneur.2016.00243

Gupta, M. D., Thakurta, R., & Basu, A. (2025). Relationship between Laterality and Theory of Mind among Typical Adults – A Systematic Literature Review. Acta Psychologica, 254, 104862. 10.1016/j.actpsy.2025.104862

Hamasaki, T., Leingärtner, A., Ringstedt, T., & O’Leary, D. D. M. (2004). EMX2 Regulates Sizes and Positioning of the Primary Sensory and Motor Areas in Neocortex by Direct Specification of Cortical Progenitors. Neuron, 43(3), 359–372. 10.1016/j.neuron.2004.07.016

Hiersche, K. J., Schettini, E., Li, J., & Saygin, Z. M. (2024). Functional dissociation of the language network and other cognition in early childhood. Human Brain Mapping, 45(9), e26757. 10.1002/hbm.26757

Hoerl, A. E., & Kennard, R. W. (1970). Ridge Regression: Applications to Nonorthogonal Problems. Technometrics, 12(1), 69–82. 10.2307/1267352

Hu, D., Wang, F., Zhang, H., Wu, Z., Zhou, Z., Li, G., Wang, L., Lin, W., Li, G., & UNC/UMN Baby Connectome Project Consortium. (2022). Existence of Functional Connectome Fingerprint during Infancy and Its Stability over Months. The Journal of Neuroscience, 42(3), 377–389. 10.1523/JNEUROSCI.0480-21.2021

Ji, J. L., Spronk, M., Kulkarni, K., Repovš, G., Anticevic, A., & Cole, M. W. (2019). Mapping the human brain’s cortical-subcortical functional network organization. NeuroImage, 185, 35–57. 10.1016/j.neuroimage.2018.10.006

Johnson, M. H. (2011). Interactive Specialization: A domain-general framework for human functional brain development? Developmental Cognitive Neuroscience, 1(1), 7–21. 10.1016/j.dcn.2010.07.003

Kanwisher, N. (2010). Functional specificity in the human brain: A window into the functional architecture of the mind. Proceedings of the National Academy of Sciences, 107(25), 11163–11170. 10.1073/pnas.1005062107

Kelly, C., & Castellanos, F. X. (2014). Strengthening Connections: Functional Connectivity and Brain Plasticity. Neuropsychology Review, 24(1), 63–76. 10.1007/s11065-014-9252-y

Khodaei, M., Laurienti, P. J., Dagenbach, D., & Simpson, S. L. (2023). Brain working memory network indices as landmarks of intelligence. Neuroimage: Reports, 3(2), 100165. 10.1016/j.ynirp.2023.100165

Li, J., Osher, D. E., Hansen, H. A., & Saygin, Z. M. (2020). Innate connectivity patterns drive the development of the visual word form area. Scientific Reports, 10(1), 18039. 10.1038/s41598-020-75015-7

Malik-Moraleda, S., Ayyash, D., Gallée, J., Affourtit, J., Hoffmann, M., Mineroff, Z., Jouravlev, O., & Fedorenko, E. (2021). *The universal language network: A cross-linguistic investigation spanning 45 languages and 12 language families* [Preprint]. Neuroscience. 10.1101/2021.07.28.454040

Marcus, D., Harwell, J., Olsen, T., Hodge, M., Glasser, M., Prior, F., Jenkinson, M., Laumann, T., Curtiss, S., & Van Essen, D. (2011). Informatics and Data Mining Tools and Strategies for the Human Connectome Project. Frontiers in Neuroinformatics, 5. 10.3389/fninf.2011.00004

McCarthy, G., Puce, A., Gore, J. C., & Allison, T. (1997). Face-Specific Processing in the Human Fusiform Gyrus. Journal of Cognitive Neuroscience, 9(5), 605–610. 10.1162/jocn.1997.9.5.605

Mennes, M., Kelly, C., Zuo, X.-N., Di Martino, A., Biswal, B. B., Castellanos, F. X., & Milham, M. P. (2010). Inter-individual differences in resting-state functional connectivity predict task-induced BOLD activity. NeuroImage, 50(4), 1690–1701. 10.1016/j.neuroimage.2010.01.002

Molloy, M. F., & Osher, D. E. (2023). A personalized cortical atlas for functional regions of interest. Journal of Neurophysiology, 130(5), 1067–1080. 10.1152/jn.00108.2023

Molloy, M. F., Saygin, Z. M., & Osher, D. E. (2024). Predicting high-level visual areas in the absence of task fMRI. Scientific Reports, 14(1), 11376. 10.1038/s41598-024-62098-9

Mueller, S., Wang, D., Fox, M. D., Yeo, B. T. T., Sepulcre, J., Sabuncu, M. R., Shafee, R., Lu, J., & Liu, H. (2013). Individual Variability in Functional Connectivity Architecture of the Human Brain. Neuron, 77(3), 586–595. 10.1016/j.neuron.2012.12.028

Ojemann, G. (1991). Cortical organization of language. The Journal of Neuroscience, 11(8), 2281–2287. 10.1523/JNEUROSCI.11-08-02281.1991

O’Leary, D. D. M., Chou, S.-J., & Sahara, S. (2007). Area Patterning of the Mammalian Cortex. Neuron, 56(2), 252–269. 10.1016/j.neuron.2007.10.010

Olulade, O. A., Seydell-Greenwald, A., Chambers, C. E., Turkeltaub, P. E., Dromerick, A. W., Berl, M. M., Gaillard, W. D., & Newport, E. L. (2020). The neural basis of language development: Changes in lateralization over age. Proceedings of the National Academy of Sciences, 117(38), 23477–23483. 10.1073/pnas.1905590117

Osher, D. E., Brissenden, J. A., & Somers, D. C. (2019). Predicting an individual’s dorsal attention network activity from functional connectivity fingerprints. Journal of Neurophysiology, 122(1), 232–240. 10.1152/jn.00174.2019

Osher, D. E., Saxe, R. R., Koldewyn, K., Gabrieli, J. D. E., Kanwisher, N., & Saygin, Z. M. (2016). Structural Connectivity Fingerprints Predict Cortical Selectivity for Multiple Visual Categories across Cortex. Cerebral Cortex, 26(4), 1668–1683. 10.1093/cercor/bhu303

Passingham, R. E., Stephan, K. E., & Kötter, R. (2002). The anatomical basis of functional localization in the cortex. Nature Reviews Neuroscience, 3(8), 606–616. 10.1038/nrn893

Rajimehr, R., Firoozi, A., Rafipoor, H., Abbasi, N., & Duncan, J. (2022). Complementary hemispheric lateralization of language and social processing in the human brain. Cell Reports, 41(6), 111617. 10.1016/j.celrep.2022.111617

Rossion, B., Hanseeuw, B., & Dricot, L. (2012). Defining face perception areas in the human brain: A large-scale factorial fMRI face localizer analysis. Brain and Cognition, 79(2), 138–157. 10.1016/j.bandc.2012.01.001

Sala-Llonch, R., Bartrés-Faz, D., & Junqué, C. (2015). Reorganization of brain networks in aging: A review of functional connectivity studies. Frontiers in Psychology, 6. 10.3389/fpsyg.2015.00663

Saygin, Z. M., Osher, D. E., Koldewyn, K., Reynolds, G., Gabrieli, J. D. E., & Saxe, R. R. (2012). Anatomical connectivity patterns predict face selectivity in the fusiform gyrus. Nature Neuroscience, 15(2), Article 2. 10.1038/nn.3001

Saygin, Z. M., Osher, D. E., Norton, E. S., Youssoufian, D. A., Beach, S. D., Feather, J., Gaab, N., Gabrieli, J. D. E., & Kanwisher, N. (2016). Connectivity precedes function in the development of the visual word form area. Nature Neuroscience, 19(9), 1250–1255. 10.1038/nn.4354

Sherman, L. E., Rudie, J. D., Pfeifer, J. H., Masten, C. L., McNealy, K., & Dapretto, M. (2014). Development of the Default Mode and Central Executive Networks across early adolescence: A longitudinal study. Developmental Cognitive Neuroscience, 10, 148–159. 10.1016/j.dcn.2014.08.002

Shine, J. M., Bissett, P. G., Bell, P. T., Koyejo, O., Balsters, J. H., Gorgolewski, K. J., Moodie, C. A., & Poldrack, R. A. (2016). The Dynamics of Functional Brain Networks: Integrated Network States during Cognitive Task Performance. Neuron, 92(2), 544–554. 10.1016/j.neuron.2016.09.018

Sporns, O. (2013). Structure and function of complex brain networks. Dialogues in Clinical Neuroscience, 15(3), 247–262. 10.31887/DCNS.2013.15.3/osporns

Squire, L. R., Stark, C. E. L., & Clark, R. E. (2004). The medial temporal lobe. Annual Review of Neuroscience, 27, 279–306. 10.1146/annurev.neuro.27.070203.144130

Sun, F., Yang, T., Liu, N., & Wan, X. (2023). The Causal Role of Temporoparietal Junction in Mediating Self–Other Mergence during Mentalizing. Journal of Neuroscience, 43(49), 8442–8455. 10.1523/JNEUROSCI.1026-23.2023

Tavoni, G., Ferrari, U., Battaglia, F. P., Cocco, S., & Monasson, R. (2017). Functional coupling networks inferred from prefrontal cortex activity show experience-related effective plasticity. Network Neuroscience, 1(3), 275–301. 10.1162/NETN_a_00014

Tavor, I., Jones, O. P., Mars, R. B., Smith, S. M., Behrens, T. E., & Jbabdi, S. (2016). Task-free MRI predicts individual differences in brain activity during task performance. Science, 352(6282), 216–220. 10.1126/science.aad8127

Thomas Yeo, B. T., Krienen, F. M., Sepulcre, J., Sabuncu, M. R., Lashkari, D., Hollinshead, M., Roffman, J. L., Smoller, J. W., Zöllei, L., Polimeni, J. R., Fischl, B., Liu, H., & Buckner, R. L. (2011). The organization of the human cerebral cortex estimated by intrinsic functional connectivity. Journal of Neurophysiology, 106(3), 1125–1165. 10.1152/jn.00338.2011

Thomason, M. E., Dassanayake, M. T., Shen, S., Katkuri, Y., Alexis, M., Anderson, A. L., Yeo, L., Mody, S., Hernandez-Andrade, E., Hassan, S. S., Studholme, C., Jeong, J.-W., & Romero, R. (2013). Cross-Hemispheric Functional Connectivity in the Human Fetal Brain. Science Translational Medicine, 5(173), 173ra24–173ra24. 10.1126/scitranslmed.3004978

Tik, N., Gal, S., Madar, A., Ben-David, T., Bernstein-Eliav, M., & Tavor, I. (2023). Generalizing prediction of task-evoked brain activity across datasets and populations. NeuroImage, 276, 120213. 10.1016/j.neuroimage.2023.120213

Tobyne, S. M., Somers, D. C., Brissenden, J. A., Michalka, S. W., Noyce, A. L., & Osher, D. E. (2018). Prediction of individualized task activation in sensory modality-selective frontal cortex with ‘connectome fingerprinting.’ NeuroImage, 183, 173–185. 10.1016/j.neuroimage.2018.08.007

Valk, S. L., Xu, T., Paquola, C., Park, B., Bethlehem, R. A. I., Vos de Wael, R., Royer, J., Masouleh, S. K., Bayrak, Ş., Kochunov, P., Yeo, B. T. T., Margulies, D., Smallwood, J., Eickhoff, S. B., & Bernhardt, B. C. (2022). Genetic and phylogenetic uncoupling of structure and function in human transmodal cortex. Nature Communications, 13(1), 2341. 10.1038/s41467-022-29886-1

Van Essen, D. C., Smith, S. M., Barch, D. M., Behrens, T. E. J., Yacoub, E., & Ugurbil, K. (2013). The WU-Minn Human Connectome Project: An overview. NeuroImage, 80, 62–79. 10.1016/j.neuroimage.2013.05.041

Yarkoni, T., Poldrack, R. A., Nichols, T. E., Van Essen, D. C., & Wager, T. D. (2011). Large-scale automated synthesis of human functional neuroimaging data. Nature Methods, 8(8), 665–670. 10.1038/nmeth.1635

Zhang, H.-Y., Wang, S.-J., Liu, B., Ma, Z.-L., Yang, M., Zhang, Z.-J., & Teng, G.-J. (2010). Resting Brain Connectivity: Changes during the Progress of Alzheimer Disease. Radiology, 256(2), 598–606. 10.1148/radiol.10091701

